# Reinforcement Expectation in the Honey Bee (*Apis mellifera*): Can Downshifts in Reinforcement Show Conditioned Inhibition?

**DOI:** 10.1101/2024.01.03.574098

**Authors:** Shawn Mahoney, Jay Hosler, Brian H Smith

## Abstract

When animals learn the association of a Conditioned Stimulus with an Unconditioned Stimulus, later presentation of the CS invokes a representation of the US. When the expected US fails to occur, theoretical accounts predict that conditioned inhibition can accrue to any other stimuli that are associated with this change in the US. Empirical work with mammals has confirmed the existence of conditioned inhibition. But the way it is manifested, the conditions that produce it, and determining whether it is the opposite of excitatory conditioning, are important considerations. Invertebrates can make valuable contributions to this literature because of the well-established conditioning protocols and access to the central nervous system for studying neural underpinnings of behavior. Nevertheless, while conditioned inhibition has been reported, it has yet to be thoroughly investigated in invertebrates. Here we evaluate the role of the unconditioned stimulus (US) in producing conditioned inhibition by using Proboscis Extension Response conditioning of the honey bee (*Apis mellifera*). Specifically, using variations of a ‘feature-negative’ experimental design, we employ downshifts in US intensity relative to US intensity used during initial excitatory conditioning, to show that an odorant in an odor-odor mixture can become a conditioned inhibitor. We argue that some alternative interpretations to conditioned inhibition are unlikely. However, we show variation across individuals in how strongly they show Conditioned Inhibition, with some individuals possibly revealing a different means of learning about changes in reinforcement. We discuss how resolution of these differences is needed to fully understand whether and how Conditioned Inhibition is manifested in the honey bee, and whether it can be extended to investigate how it is encoded in the CNS. It is also important for extension to other insect models. In particular, work like this will be important as more is revealed of the complexity of the insect brain from connectome projects.

## INTRODUCTION

Animal studies of Pavlovian and operant conditioning have revealed important, generalizable principles of reinforcement learning(Sutton and Barto, 2018). Many studies have now shown that animals assess the informative value of stimuli and then update behavioral decisions based on that assessment every time these stimuli occur. At its most fundamental level, associative (Pavlovian) conditioning involves pairing of a ‘neutral’ Conditioned Stimulus (CS) with an important, salient Unconditioned Stimulus (US), such as food as an appetitive example or painful stimuli as aversive examples(Rescorla, 1988). The predictive nature of a CS produces a conditioned behavior that prepares animals for the encounter with the US. These principles have been expressed in simple mathematical equations aimed at explaining how conditioning establishes associative strength V_A_ to a CS_A_, which then is the basis for anticipatory behavior(Bush and Mosteller, 1951; Rescorla and Wagner, 1972). Although there are important differences between the different models, such as whether and how attention contributes to conditioning(Lubow, 1989; Mackintosh, 1975), these mathematical formulations share two important principles(Pearce and Hall, 1980; Rescorla, 1988). First, the equations assume that the change in associative strength ΔV can be either positive when the CS predicts the US will occur, or under at least some conditions negative when the CS predicts that an expected US will *not-occur*. Second, they all propose that ΔV on a given trial is proportional to the maximum amount of associative strength that a US can support (λ) minus an expectation of the summed total associative strength to all stimuli present – including contextual stimuli - that has already accrued regarding that US (V_Σ_=V_A_+V_B_+…V_N_). Thus, associative strength V increases with each trial as long as λ>V_Σ_, i.e. as long as the US that is encountered is stronger than predicted by V_Σ_ (a positive prediction error). Once the US is fully predicted (λ=V_Σ_) associative strength stops being incremented, resulting in an asymptotic acquisition curve.

Behavioral studies have identified important manifestations of *reinforcement expectation* based on this type of model(Dunsmoor et al., 2015; Pearce and Hall, 1980; Rescorla and Wagner, 1972; Sutton and Barto, 2018). When an animal experiences CS_A_ after it has been associated with reinforcement, CS_A_ evokes a prediction of the US that is proportional to V_Σ_. This amounts to an expectation that a US at least of value V_Σ_ will occur. When a second CS_X_ is combined with CS_A_, and the same reinforcement is presented, animals typically learn less about CS_X_ than they normally would, which is called *blocking(Mackintosh, 1983)*. Additionally, it is important to establish what happens on trials when the US fails to occur despite being predicted by CS_A_, that is, when there is a surprising reward *omission(Papini, 2003)*. In particular, when a second CS_X_ is present in compound (i.e. CS_AX_) during reward omission, *conditioned inhibition to CS*_*X*_ may ensue (Rescorla, 1969). Essentially, on such CS_AX_ trials λ=0 because the US is absent. However, V_Σ_>0 because of the presence of CS_A_. Therefore λ-V_Σ_ is negative, reflecting that the US the animals encounter is less strong than what they expected. This *negative prediction error* serves as a teaching signal to establish a negative prediction for CS_X_, i.e., the prediction that the US will not-occur. This is why CS_X_ becomes a *conditioned inhibitor*.

Revealing whether and under what conditions an animal develops and expresses Conditioned Inhibition can be critical for understanding how that animal is capable of representing information about reinforcement, and in particular which of the several models of conditioning apply (e.g. whether there is a representation akin to V_Σ_ or whether CS_AX_ is perceived as a different stimulus altogether). Moreover, showing the existence of Conditioned Inhibition will be important for understanding how that capability is supported by neural connectivity in the brain.

However, it is not trivial to unambiguously demonstrate Conditioned Inhibition (Papini and Bitterman, 1993; Sosa and Ramírez, 2019). Minimally, a conditioned inhibitor should pass two types of tests (Rescorla, 1969). First, once a stimulus (CS_X_) has taken on inhibitory properties, subsequent excitatory conditioning should proceed more slowly than it does with other stimuli that do not carry inhibitory properties. A lack of retardation, or even its opposite, better excitatory conditioning of CS_X_, might indicate that more attention is devoted to the processing of CS_X_ at the expense of other stimuli that are present. Second, CS_X_ should pass a summation test. When CS_X_ is presented in compound with a second CS that itself is capable of releasing a response, that response to the compound should be diminished by the inhibitory properties of CS_X_.

The first step in this process is to demonstrate behavior that is consistent with Conditioned Inhibition. Although the existence of reward expectations has been well documented in vertebrates (Sosa and Ramírez, 2019), it has not been well studied via retardation and summation tests in invertebrates, in spite of their widespread use as models for reinforcement learning. Using the honey bee (*Apis mellifera*), Chandra (Chandra et al., 2010) showed that unreinforced presentation of an odor X – as it would occur in an AX compound without reinforcement but where A predicts reinforcement, reliably produces retardation of acquisition. In other experiments, a reduction in expected reinforcement reduced learning of a target CS below that expected given the level of reinforcement(Smith, 1997). Furthermore, backward pairing (Hellstern et al., 1998) and peak shifts (Fernandez et al., 2009) produce inhibition-like effects. These experiments seem to rule out attention as a factor. However, the focus of these works was not on Conditioned Inhibition, so the results of summation tests were not evaluated.

Here we use variants of a feature-negative (A+|AX-) experimental design to test more directly for evidence of negative summation consistent with conditioned inhibition in the honey bee PER procedure. We show that when associated with omission of expected reinforcement, X can reduce the response to another excitatory stimulus. We also show and discuss that individuals differ in expression of Conditioned Inhibition, which parallels other studies of individual differences in learning performance in honey bees, and it brings into focus how potentially multiple conditioning phenomena can affect behavior. Resolution of these differences will be necessary to more fully reveal and prove the existence of Conditioned Inhibition, which can also serve as a model for extension of these kinds of studies to other insects. Finally, use of more complex and well controlled conditioning protocols to study learning should help map these phenomena onto the unexpected complexity of the fruit fly brain connectome(Li et al., 2020), and in particular it may reveal whether and how structures like the mushroom bodies are involved in reward expectation.

## RESULTS

### Experiment 1: First test for negative summation

Subjects were initially conditioned in two phases. First, they were conditioned to A++ and B+ on separate trials that were pseudorandomly interspersed with one another (Fig. 1A solid lines and filled symbols). For the second phase, subjects were conditioned to AX+ on trials that were similarly interspersed with continued presentation of A++ (Fig. 1A dashed lines and open symbols). The AX+ condition was designed to build inhibitory tendencies to X because of the decrease in reinforcement from A++ to AX+.

**Figure 1.**
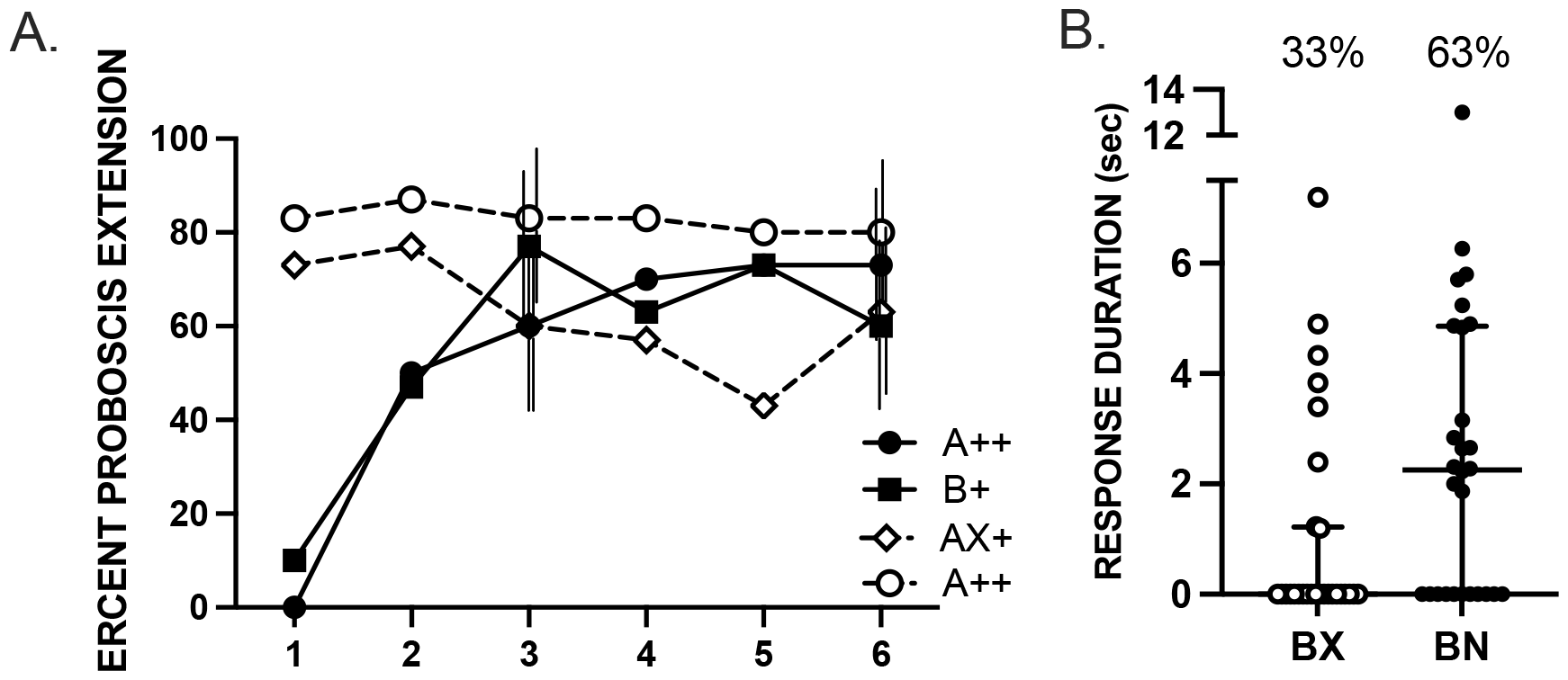
Summation test for conditioned inhibition. A, B, N and X were the odor Conditioned stimuli (CS). (A) Subjects (n = 28) were preconditioned over 12 conditioning trials (solid lines, filled symbols). Six were performed with A++ and 6 with B+ presented in a pseudorandomized sequence. In the compound conditioning phase (dashed lines and open symbols), subjects were trained over six training trials each to A++ and AX+. Vertical lines for trials 3 and 6 indicate 95% confidence intervals offset left to right for A++, B+, AX+ and A++. See Methods for calculation and limits to comparison of different treatments. (B) In the within-subjects design, all subjects were tested with the odor mixtures BN and BX. Horizontal lines represent the medians for each group, and vertical capped lines interquartile ranges. Symbols are the durations for individual subjects. Percentages about each set of points indicate the percentage of bees that responded in each group.

There was little difference between the preconditioning acquisition curves for odor A and odor B (Fig. 1A). In the compound conditioning phase, response to odor A alone (reinforced with High sucrose) remained high while the percent proboscis extension in the AX+ group (AX reinforced with Low sucrose) trended toward an overall decline. Since the strength of the US associated with A decreased between the preconditioning and compound conditioning phases in the AX+ trials, the decline in proboscis extension may already suggest an accrual of inhibition to odor X. If inhibition has accrued to X in the compound phase as a result of its association with a lower-than-expected US intensity, then this inhibitory effect should influence the subjects’ relative response to the otherwise excitatory odor B when it is paired with X during testing.

We then tested subjects with a blend of B and X. Under normal circumstances, the addition of an odorant to a previously conditioned B would decrease the response to B because of overshadowing or interactions in sensory transduction processes(Smith, 1998). Therefore, we compare the response duration to B when it is combined with X to the response duration to B when it is combined with a novel odor (N). The inhibitory properties of X should decrease the response to B more than the addition of a novel odor N that is neutral.

In the final summation test, the response to odor B should be suppressed in the BX mixture (because of its pairing with putative inhibitor X) relative to its response to odor B when it is paired to the novel odor N. The results indicate that this is indeed the case (Fig. 1B). The duration of proboscis extension response to BX was significantly shorter than that to BN (W = 101.0, p<0.01 one tailed test). The distributions shown in Fig. 1B reflects a number of subjects that failed to respond to one or both odors, which are the points lying on the x-axis.

### Experiment 2: A feature-negative test of Conditioned Inhibition

Most studies of conditioned inhibition have used a variation of a feature-negative design(Papini and Bitterman, 1993), which is a somewhat different procedure from that in our first experiment. In a feature negative experiment we employ here, one odorant (A++) is paired with reinforcement, whereas a mixture of that odorant with a target odorant (AX-) is not reinforced at all. We have used a similar procedure to demonstrate feature-negative conditioning in the honey bee(Chandra and Smith, 1998). This procedure produces a much more dramatic discrepancy between the reinforcement signaled by A and the actual outcome on trials when X is present. As in the first experiment, we used a single treatment group of 24 subjects. Each subject experienced four different types of trials that were presented intermixed in a pseudorandom order. Two types of trials involved pairing an odorant with either ++ or + reinforcement (A++ and B+) on separate trials. In this experiment we used B as an excitatory target odor against which to test for a summation effect. There were also two types of completely unreinforced trials (AX- and CY-). We predicted that inhibition would accrue to X on AX-trials because of the discrepancy between the signaled and actual outcomes. Furthermore, because unreinforced presentation of a CS may reduce its potential excitatory properties via latent inhibition(Chandra et al., 2010), we present a second unreinforced odor blend (CY-). If the inhibition that accrues to X is simply due to attentional decrement from nonreinforcement (latent inhibition), then there should be no difference between the response levels BX and BY in a subsequent summation test. On the other hand, if the lack of reinforcement in an otherwise excitatory context produces conditioned inhibition, then we would expect that the response to BX would be less than that to BY. Furthermore, if afferent interaction in the AX compound reduces the response to BX, then there should be an equivalent reduction in BY because of afferent interaction in the CY condition.

Subjects showed rapid acquisition to A++ and to B+ (Fig. 2A), reaching an asymptotic level of response (circa 70%) by trials 4 or 5. The response level to AX-was intermediate, reaching a final response level of circa 25%. The response to CY-, the components of which were never reinforced, was lowest. The somewhat elevated level of response to AX- as compared to CY-could be due to incomplete excitatory generalization from A to the AX compound. Furthermore, inhibition to X could have also contributed to the response to AX-in the sense that it would decrease the response to AX relative to A.

**Figure 2.**
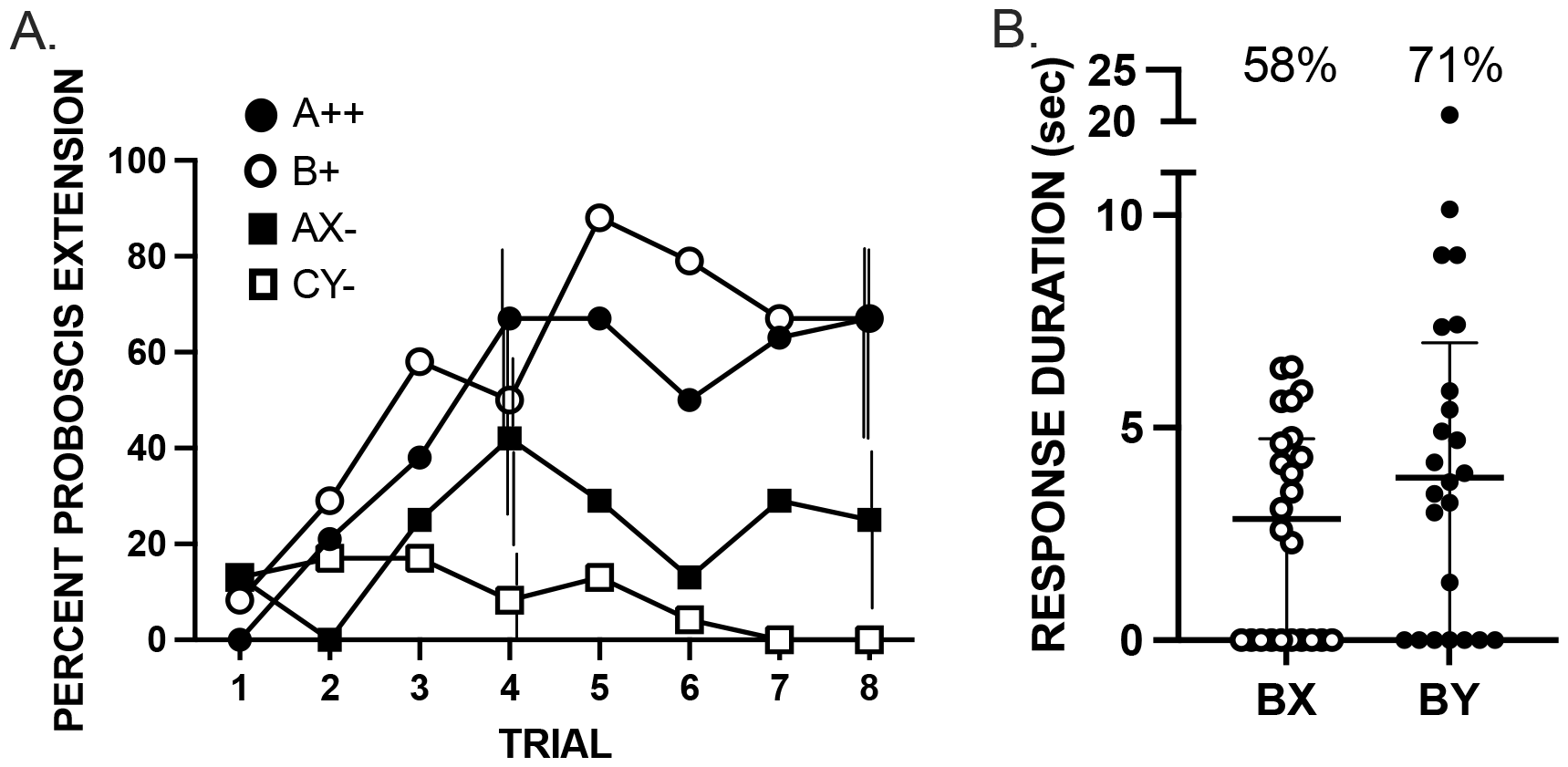
Test for conditioned inhibition using a feature-negative design. All subjects (n = 24) were exposed to 8 conditioning trials in each of the following 4 treatments: A++/B+/AX-/CY-. The 4 odorants used in this experiment were counterbalanced as A, X, C and Y. B was an excitatory test stimulus, and was always geraniol. (A) Acquisition to all 4 treatments. Vertical lines for trials 3 and 6 indicate 95% confidence intervals offset left to right for A++, B+, AX- and CY-. See Methods for calculation and limits to comparison of different treatments. (B) Unreinforced summation tests with BX and BY. Horizontal lines represent the medians for each group, and vertical capped lines interquartile ranges. Symbols are the durations for individual subjects. Percentages about each set of points indicate the percentage of bees that responded in each group.

Indeed, the summation test reveals that there is most likely a contribution from conditioned inhibition. The response to BX was significantly less than that to BY (Fig. 2B; W = 102.0, p<0.05 one-tailed test). Thus, odor X was capable of more significant negative summation when added to the conditioned excitor B than when Y was added.

### Experiment 3: Individual differences in expression of Conditioned Inhibition

Figures 1 and 2 reflect mean test performance across a sample of 24 or 30 subjects, which is standard procedure for presenting learning performance in PER studies(Burden et al., 2016). However, mean values hide diversity of performance on any learning task, with some subjects performing well and others not showing the expected outcome. These differences are reflected in the distributions of individual duration values shown in Figs. 1B and 2B. Because of these individual differences, and its importance for a social colony (see Discussion;(Cook et al., 2019)), we decided to repeat the second experiment (Fig 2) with a focus on individual differences.

Acquisition across the four odors in this replicate were similar to acquisition in the first experiment (Figs. 2A). The response levels across trials to reinforced odors A++ and B+ were highest, reaching 70 to 80% of subjects responding or higher by the 5^th^ or 6^th^ trials. As before the response level to AX-was intermediate and the response to CY-was lowest.

In this replicate, differences between BX and BY failed to reach significance, even though the trend was in the same direction as in experiment 2 (median BX 2.7 vs BY 3.2). The differences between experiments are a major point in our discussion below. In the meantime, some important clarification can be provided by a more detailed inspection of the data in both experiments.

In both experiments a majority of subjects that responded to one or both odors (13 of 20 and 19 of 32, resp) responded more strongly – i.e. with longer response durations - to BY than to BX (Fig. 4A,B; *X*^*2*^ p<0.01), which is the pattern expected for Conditioned Inhibition. Data from these subjects are plotted as ‘Inhibitors’. However, it is notable that several subjects in each experiment failed to show the pattern predicted by the Conditioned Inhibition hypothesis. First, some individuals failed to respond to either odor compound (4 and 9 in experiments 2 and 3, resp). Second, some subjects (7 of 20 and 13 of 32 in experiments 2 and 3, resp) showed shorter response durations to BY than to BX (right pairs of columns labelled ‘Non-inhibitor’), which is opposite to the prediction for Conditioned Inhibition. In fact, in both experiments some subjects (shaded symbols) responded to BX but failed to respond to BY. Obviously, because of the way they were selected, the differences within each group are statistically significant (p<0.01). The implications of these differences are further discussed below.

**Figure 3.**
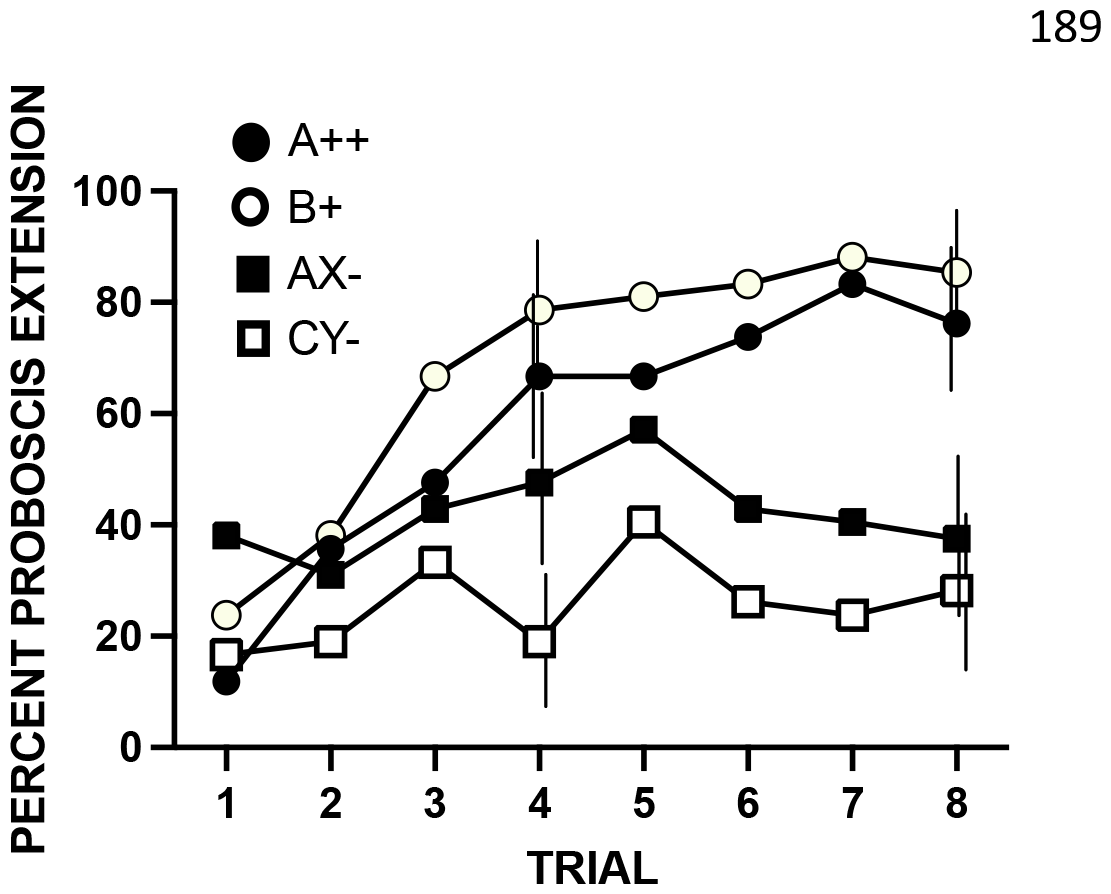
Acquisition phase for the replicate of experiment 2 using a feature-negative design. All subjects (n = 41) were exposed to 8 conditioning trials in each of the following 4 treatments: A++/B+/AX-CY-. The 4 odorants used in this experiment were counterbalanced s A, X, C and Y. B was an excitatory test stimulus, and was always geraniol. Vertical lines for trials 3 and 6 indicate 95% confidence intervals offset left to right for A++, B+, AX- and CY-. See Methods for calculation and limits to comparison of different treatments.

**Figure 4.**
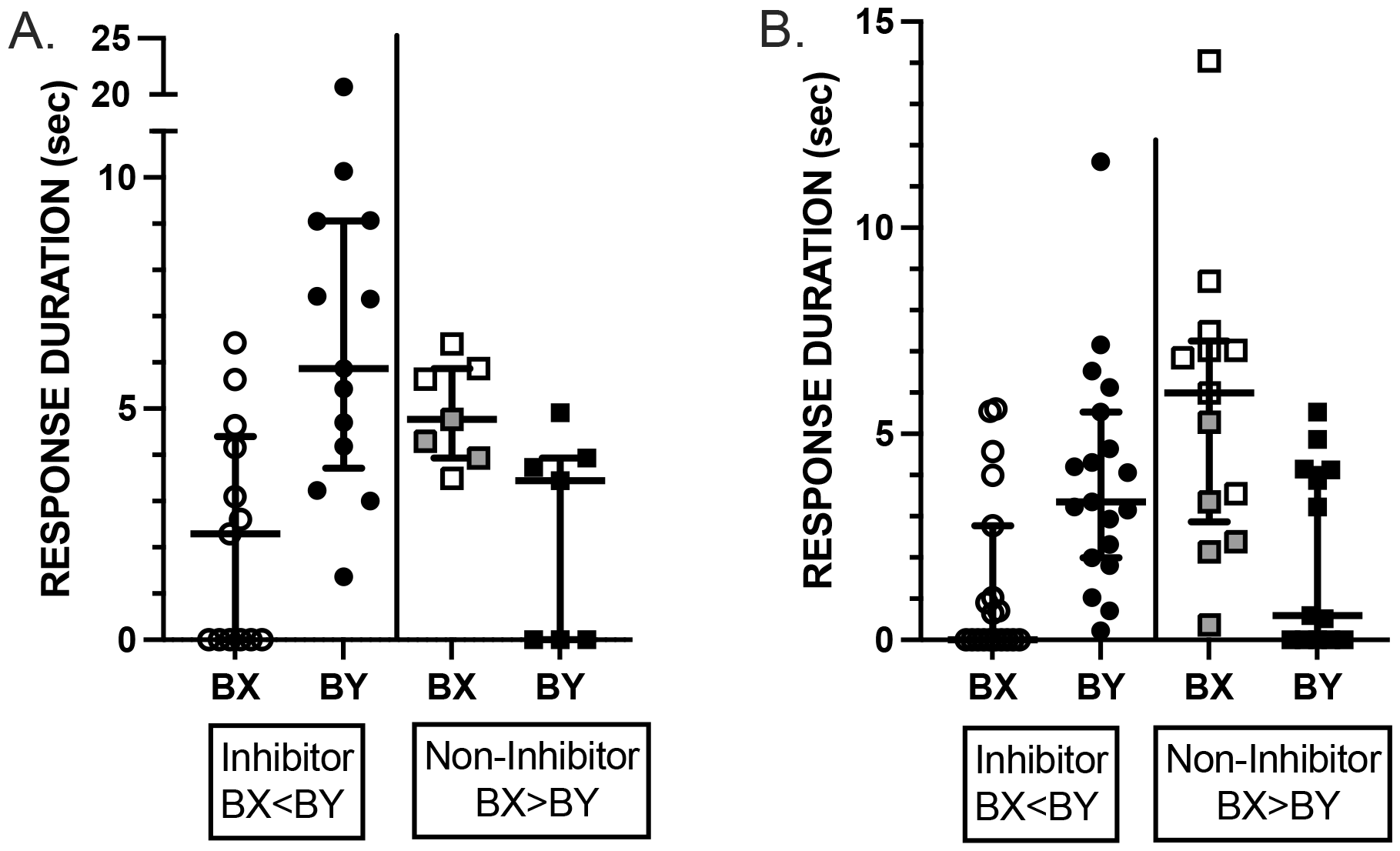
Summation tests for experiments 2 (A) and 3 (B). Plots show data from subjects that responded to one or both odors in unreinforced summation tests with BX and BY. Subjects that failed to respond to either odor (4 in A and 9 in B) are not shown. Subjects in each experiment were classified as ‘Inhibitor’ if they showed a stronger response to BY than to BX. Otherwise, they were classified as Non-Inhibitor. Horizontal lines represent the medians for each group, and vertical capped lines interquartile ranges. Symbols are the durations for individual subjects. Percentages about each set of points indicate the percentage of bees that responded in each group. Shaded symbols for Non-Inhibitors are for subjects that responded to BX but not to BY. See text for proportion data.

## DISCUSSION

Our results suggest that a component of a binary odor blend can develop properties of a conditioned inhibitor (Papini, 2003; Rescorla, 1969; Sosa and Ramírez, 2019). These properties develop when that component is associated with a lower level of reinforcement than that signaled by other components of the blend. However, as we discuss below, we must use caution at this point in reaching firm conclusions because of individual variation that we report. In addition, there are several alternative interpretations that can account for a decrease in response to a target stimulus(Papini and Bitterman, 1993; Sosa and Ramírez, 2019). Those alternatives include modification of attention, afferent interaction, external inhibition, changes in generalized excitation, US habituation and contextual conditioning/blocking. It is important now to address these issues in order to determine the reliability of the claim for inhibition in our experiments.

Rescorla (Rescorla, 1969) has recommended a two-test strategy for revealing conditioned inhibition, which would serve as a necessary set of conditions for showing inhibition. A conditioned inhibitor would reveal retardation of acquisition when it is properly paired with reinforcement, and its inhibition would negatively summate with the excitatory properties of another stimulus that was associated with the same type of reinforcement used to produce the inhibition. If A+/AX-pairing changed attention to X, then X might pass one test but not the other. For example, if attention to X were enhanced because of the *surprising* omission of reinforcement in the A+/AX-condition, then acquisition would be expected to be more rapid to X in an A+/AX-group than in a control group (e.g., A+/X-). In this case X would fail the retardation test but might still decrease the response to an excitatory stimulus because of X’s ability to monopolize attention. On the other hand, if the effect of nonreinforcement of X in the A+/AX-condition were to decrease attention to X via latent inhibition(Lubow, 1973), for example, then acquisition would be retarded, and X would pass the retardation test, but X might fail to reduce the excitatory properties of another stimulus leading to failure of the summation test. Only a conditioned inhibitor would be capable of passing both tests.

Interpretation of reduced response to BX in our experiments, relative to BN, because of increased attention to X is unlikely. If attention were a factor, then any modification of the US between the initial and compound trials will generate some form of surprise and hence enhanced attention to X(Mackintosh, 1983; Pearce and Hall, 1980). That is, either an up- or a down-shift in reinforcement should be capable of generating a significant amount of learning about X on compound (AX) trials. However, when honey bees experience A++/AX++ training, they learn less about X relative to a control procedure in which A is not present during the compound phase(Smith, 1997). Furthermore, when they experience a downshift in reinforcement (A++/AX+), they respond less to X relative to the A++/AX+ group that did not experience the downshift. The attention hypothesis would predict the opposite. Moreover, this procedure showed one-trial blocking. Those results are inconsistent with the interpretation that attention to X is enhanced by a downshift in reinforcement used in our current experiments(Mackintosh, 1983; Rescorla and Holland, 1982).

Latent inhibition (Lubow, 1973) via retardation of acquisition also does not account for our results. Latent inhibition in the honey bee PER paradigm requires many more (∼20) unreinforced exposures to a CS (Chandra et al., 2010) than subjects were exposed to in our experiment. Nevertheless, we attempted to control for this effect in experiments 2 and 3 by exposing subjects to the CY-condition. If latent inhibition decreased response to B in the AB-condition, then it should have done so equally to Y in the CY-condition. But the test response to BX was lower than to BY, which argues at least for an additional source of inhibition to X.

Afferent interaction might reasonably be expected to be stronger in intramodal compounds than in intermodal compounds(Kehoe et al., 1994). We attempted to control for afferent interaction by the addition of a novel odorant during test phase, as we previously have done for studies of blocking(Smith, 1996; Smith, 1997; Smith and Cobey, 1994; Thorn and Smith, 1997). If we had simply tested BX versus B, then the decrease would be interpretable in terms of greater interaction in the compound, which occurs in mixtures of these same odorants used above(Smith, 1998). Furthermore, we either partially or completely counterbalanced odorants in all of our experiments. It is therefore unlikely that the summation effect of X would arise due to greater afferent interaction with B than in the control groups.

Several previous attempts have been made to specifically study conditioned inhibition in honey bees, but the results were mixed. Some early experiments using different types of manipulations(Bitterman et al., 1983), including backward pairing (Hellstern et al., 1998) and peak shift(Fernandez et al., 2009), provided an indication of inhibition, but the experiments left open alternative interpretations. Couvillon et al. (Couvillon et al., 1999)failed to find evidence for conditioned inhibition in freely flying honey bees. Yet, use of modified, and perhaps more sensitive, experimental designs detected evidence of conditioned inhibition(Couvillon et al., 2003). Using restrained subjects in the PER paradigm provided indirect evidence for inhibition(Couvillon et al., 2005). Yet our experiments using PER reported here support the interpretation, over other alternative possibilities, that conditioned inhibition might still be found in honey bees. The design we adopted for experiments 2 and 3 was adopted from Couvillon et al.(Couvillon et al., 1999). Therefore, differences across studies do not appear to be attributable to use of freely flying versus restrained bees, or to the use of intra-versus inter-modal stimulus mixtures.

A major difference between our study reported here and, other studies using PER conditioning of restrained subjects (Couvillon et al., 2005) involves aspects of experimental design and use of different response measures. Our designs employed longer phases of excitatory conditioning – 18 or 32 trials versus 9. A similar difference can account for differences in detection of latent inhibition(Bitterman et al., 1983; Chandra et al., 2010). Furthermore, studies of PER conditioning have mostly used the percentage of subjects in a treatment group that responded to the CS. This measure is frequently sensitive enough to detect differences between treatment groups. But failure to find differences, particularly when a large percentage of subjects respond, cannot be interpreted as unambiguous evidence of a lack of treatment effect. Although the response itself looks simple, detailed electromyographic (Smith and Menzel, 1989) and video analyses like we have done here reveal several different features of response topology (e.g. response latency and duration) that differentiate treatment groups(Smith, 1997), even when no differences are evident in response probability.

Here we have tried to be careful to point out individual differences in manifestation of Conditioned Inhibition, which are present in all studies of PER conditioning but are often not apparent when mean values are presented. Several other studies of PER in honey bees have also shown consistent individual differences in performance in several different learning protocols (Finke et al., 2021; Finke et al., 2023; Scheiner et al., 2017; Tait et al., 2019), which also broadly correlates to sensitivity to sugar as well as to other foraging preferences (Scheiner et al., 2001). Genotype accounts for the largest proportion of variance in learning performance among bees in the same colony (Bhagavan et al., 1994). Once genotype is more standardized, differences in learning performance between behavioral castes become more apparent (Ferguson, 2001). Because a queen mates with up to 20 drones (males), any colony will contain a mixture of several different patrilines (Page, 2013). This genetic diversity within any colony is important for the fitness of the colony as it performs foraging tasks (Cook et al., 2020; Mosqueiro et al., 2017). However, this diversity must be taken into account in any study of conditioning, because a small sample (∼20 or 30) workers from a large population (up to 100,000) may not reflect the genetic diversity present in the large population. This could be the reason that it was more difficult to show in experiment 3 than in experiment 2, for example.

A few loci in the honey bee genome correlate to learning performance, but one in particular has been highly correlated in two independent genetic mapping studies (Chandra et al., 2000; Latshaw et al., 2023). In this locus, a gene that encodes a receptor for the biogenic amine tyramine has a major impact on learning performance, and because of where that receptor is expressed (Sinakevitch et al., 2017), and its intracellular signaling pathway(Blenau et al., 2000), it is likely that it acts as a gain control to regulate inputs to circuitry that supports learning (Latshaw et al., 2023). The activity of this gene, or the lack of activity, would cause bees to display strong or poor learning performance. This model is also consistent with this gene having broad pleiotropic effects on other behaviors, for example, specializations in pollen versus nectar collation(Hunt et al., 1995). Because of this broad pleiotropy, it is possible that it affects expression of conditioned inhibition.

We confirm here that the groups of subjects selected for experiments 2 and 3 show differences across individuals in how the behavior is manifested. The majority of individuals responded more strongly to the BY compound than to the BX compound, which is the prediction for Conditioned Inhibition. In contrast, a minority of individuals in each experiment responded more strongly to BX than to BY, which is the opposite prediction from Conditioned Inhibition. This pattern (BX>BY) could arise if some individuals associated X with excitatory conditioning of A++, as would be expected for Second Order conditioning(Mackintosh, 1983), for example. In fact, the procedure we have employed would also be employed to study second order conditioning. In that case, stronger response to the BX compound could occur via *positive* summation instead of the negative summation predicted for Conditioned Inhibition.

These individual differences in our studies leave open possibilities for further investigation. For example, do some individuals in a colony show excitatory second order conditioning over Conditioned Inhibition, or vice versa, when a stimulus is paired with a conditioned excitor? Does excitatory second order conditioning drive responding over the first few conditioning trials and then switch to Conditioned Inhibition after several trials? In this case, some individuals may make this switch earlier than others, which would account for individual differences in our study. Within colony genetic differences have been shown, for example, for Latent Inhibition and Reversal Learning(Chandra et al., 2010; Ferguson, 2001), and could carry over to other forms of conditioning. Our results emphasize that future studies need to describe individual differences in learning performance and test how other factors – age, experience, behavioral caste, genetic background – affect expression of the trait. At the very least, care must be taken to control for and standardize genetic background, which is not often done in studies with honey bees, but is standard practice in studies with fruit flies and rodents.

Taken together, studies now show enough evidence that conditioned inhibition can be studied in honey bees, and now also in molluscs(Acebes et al., 2012). However, much needs to be done to verify it and more fully evaluate how conditioned inhibition is specifically manifested in behavior. One important contribution of our work reported is to provide guidance on how to study it – using the protocol in experiments 2 and 3 - and raise some important questions going forward. For example, does conditioned inhibition show extinction, like in excitatory conditioning? What is the role of contextual cues in generating and releasing memory for conditioned inhibition? Are there differences in manifestation of conditioned inhibition between inter- and intra-modal mixtures, as were used in our study? As noted above, is second order conditioning expressed in the first few trials with conditioned inhibition more dominant with longer sequences of trails? Answers to these questions will be essential for understanding how different forms of learning are integrated into the ongoing decision processes involved in acquiring resources or avoiding threats(Bazhenov et al., 2013). Additionally, this information will interface with investigations into the neural mechanisms that underlie Conditioned Inhibition, such as have been initiated for latent inhibition (Locatelli et al., 2013) and blocking (Chen et al., 2015) in honey bees. These studies in insects such as honey bees and fruit flies, particularly if conditioned inhibition can be found in the latter, will help integrate the behavior into information now coming out of the fruit fly brain connectome projects(Li et al., 2020).

## METHODS

### Subjects

The subjects were honey bee workers obtained from three to five colonies maintained out-of-doors in a genetically closed population(Page et al., 1985). Subjects were collected from colony entrances, as they departed from or returned to the colony, in the morning between 9 and 10 am. They were then set-up in restraining harnesses and conditioned according to published procedures described below (and in(Smith and Burden, 2014)). After a 15 minute period to acclimate to the harness, subjects were fed 0.4µl of a 1.25M sucrose solution, and they then remained undisturbed for 2 hours before the start of the conditioning procedure. All three experiments used a within-subjects design for testing.

### Apparatus

Conditioning and testing were conducted in a 13 cm x 14.5 cm x 16.5 cm conditioning station illuminated with a fiber optic light source. The station had an open front, top and floor. The remaining sides were constructed of clear Plexiglas. To evacuate odors during conditioning, a 9.5 cm diameter hole in the back of the chamber was connected to an exhaust hood via a dryer tube. The floor of the chamber consisted of a single strip of Plexiglas connecting the left and right sides of the conditioning station. In the center of this strip was a small Plexiglas peg on which harnessed subjects were placed during conditioning.

Subjects were positioned to face an odor cartridge at the front of the chamber. Odor cartridges were made fresh daily by placing a strip of filter paper containing 3 µl of pure odorant into a glass syringe. The open tip of the odor cartridge was positioned in front of and facing the subject. The ground glass fitting was connected to a three-way valve system that regulated the flow of air being generated by an aquarium air pump. Administration of the CS (odor) was controlled by computer or by an Arduino-based logic controller. When triggered, the computer operated a solenoid valve, thus shunting air-flow away from an exhaust tube and through the odor cartridge. The odor-saturated air was ejected into the air stream that moved across the subject’s antenna. Just behind the conditioning stage and out of the subject’s view was a small LED used to signal the onset of odor delivery on videotape (see below). The US (0.4 µl of either 1.25 M or 0.50 M sucrose-water solution was administered using a Gilmont microliter syringe. The odors were hexanol, octanol, 2-hexanone, 2-octanone, geraniol and eugenol; they were rotated across days into each of the odor types (labeled as A, B, X, N, Y or C in different experiments) used in conditioning phases described below. All odors are easily discriminable to honey bees in PER conditioning.

At the start of each trial the subject was placed into the conditioning station 20 seconds prior to the onset of the odor CS. It was also left in the conditioning station 30 seconds after odor exposure. The inter-trial interval for each subject was fixed at 6 min. A video camera was positioned above the conditioning stage to record each subject’s response during the test phase of the experiment.

### Procedure

#### Proboscis extension conditioning

In this protocol(Smith and Burden, 2014), an odorant (CS) is blown across the subject’s antenna for 4 seconds. Three seconds after onset of the odor CS, a 0.4µl drop of sucrose-water (US) is lightly touched to the antenna. This stimulates sucrose taste receptors in the antenna and triggers the proboscis extension response. The subject is then allowed to consume the entire droplet, which always occurs within ca 1 sec. After 3-4 trials most subjects extend their proboscis to odor prior to sucrose reinforcement, which we registered as a positive Conditioned Response.

#### Experiment 1

The first experiment was divided into two acquisition phases during which six subjects were conditioned each day, so the intertrial interval was fixed at 6 min. In the initial (Pretraining) phase of all experiments, subjects were equivalently conditioned to six trials each of odor A and odor B presented in a pseudorandomized sequence (ABBABAABABBA). Each odor was appetitively reinforced, but with different sucrose concentrations. Odor A was reinforced with high (++; 1.25 M) sucrose and odor B was reinforced with low (+; 0.50 M) sucrose.

After pretraining, the first experiment consisted of a second acquisition phase followed by a test phase. For compound conditioning in the second phase all subjects received 12 trials. Subjects were exposed to odor A followed by High sucrose (A++ condition) on six of the trials. This condition was a continuation of the A++ condition from the preconditioning phase. On the remaining six trials, subjects were exposed to the compound of A and X followed by Low sucrose (AX+). Thus, X was reinforced with low sucrose in a context that predicted High sucrose reinforcement. The two trial types were presented in the pseudo-randomized order used for pretraining.

After the second phase each subject was tested for their response to the two odor compounds BX and BN without the presence of reinforcement. Responses to odor B in combination with odor X were compared to responses to odor B when it was mixed with the novel odor N. The odors used for A, B, X, and N, and the order of presentation of BX and BN during testing, were counterbalanced over days of the experiment.

#### Experiments 2 and 3

Each subject was exposed to 32 acquisition trials, which was comprised of 8 trials each with A++, B+, AX- and CY-in a pseudorandomized sequence across trials. The major qualitative differences between this experiment and those reported above were: (1) that trials with A++ and AX-were pseudorandomly interspersed instead of presented in successive phases; and (2) the lack of reinforcement of the AX compound instead of reinforcement with a diminished US.

Following the acquisition trials, each subject was tested on unreinforced trials with BX and with BY in a random sequence across subjects. There are 24 different combinations of 4 odorants; therefore, each of the 24 subjects was conditioned to a different pattern of A, C, X and Y. The fifth odorant (B), which was a common test stimulus, was the same for all subjects (geraniol).

### Data analysis

After conditioning was complete all subjects received a unreinforced presentation of odors BX and either BN (experiment 1) or BY (experiments 2 and 3) presented in a randomized order across days. All responses were video recorded for offline analysis. We then calculated the duration of proboscis extension during the 4 second presentation of odor X and for 20 seconds after the termination of odor presentation(Smith, 1997; Smith, 1998). Statistical analysis of the response duration was used for all hypothesis testing(Smith, 1998; Smith and Menzel, 1989). Because of the distribution of the scores, statistical analyses were accomplished with Wilcoxon signed ranks tests (Sokal and Rohlf, 1994) in PRISM (©). Medians and interquartile ranges are shown for unreinforced tests with BX, BN and BY in the figures along with symbols indicating durations from individual responses.

We present acquisition data for descriptive reasons. But we do not provide statistical tests of acquisition data, primarily because we do not use them for hypothesis testing. In particular, testing differences during acquisition trials, during which groups receive different treatment conditions, would confound treatment and testing conditions (Rescorla, 1988). For example, A++ and B+ trials presented during acquisition differ in two factors; odor and concentration of sucrose in the reward. Bees learn the visual stimulus of the water droplet approaching them and the water vapor that surrounds it. On the B+ trials, when a lower concentration is presented, bees may generalize the association of the droplet with high reward presented on the A++ trials to the B+ trials, which would elevate the response to B. In addition, the differences between rewarded trials and unreinforced (e.g. AX- or CY-) trials also involve two factors: odor and presence/absence of reward. The two-factor differences between these curves are necessary to set up different treatment conditions, but it makes them not comparable for hypothesis testing. To test for inhibition, we have added unreinforced trials at the end, which differ in only one factor (e.g. BX and BY). Furthermore, it would be impossible to calculate duration of the conditioned response when its expression would be influenced by the presentation of the US prior to offset of the CS. Given the similarity between the CR and the UR (Smith & Menzel, 1989), it would be impossible to distinguish them. We did not insert extinction trials to measure duration during acquisition, because it was not our intent to study differences in response to stimuli presented during the conditioning phases, and doing that might have attenuated the ability of a CS to induce inhibition.

Therefore, the only response measure we used for acquisition was the percentage of subjects that responded to odor prior to presentation of the CS. That measure has been used in many studies of PER conditioning in the honey bee(Menzel, 1990), and when it reveals differences across treatment groups those differences are robust with almost any measure of the CR(Smith, 1998). But when it fails to reveal differences across treatments, use of a more parametric measure such as duration or latency can reveal differences in the response topology(Smith and Menzel, 1989).

## Notes

This work was supported by awards the BHS from the National Institutes of Health (GM 113967), the National Science Foundation (2014217) NeuroNex program, the National Science Foundation (2113179) CRCNS US-France (ANR) collaborative research program, and the Department of Energy (SC0021922) US-German (BMBF) collaborative research program.

### Competing Interest Statement

The authors have declared no competing interest.

### Summary of Updates

Statistical analyses Also limitation of conclusions in light of individual variation

